# Direct detection of molecular intermediates from first-passage times

**DOI:** 10.1101/772830

**Authors:** Alice L. Thorneywork, Jannes Gladrow, Yujia Qing, Marc Rico-Pasto, Felix Ritort, Hagan Bayley, Anatoly B. Kolomeisky, Ulrich F. Keyser

## Abstract

All natural phenomena are governed by energy landscapes. However, the direct measurement of this fundamental quantity remains challenging, particularly in complex systems involving intermediate states. Here, we uncover key details of the energy landscapes that underpin a range of experimental systems through quantitative analysis of first-passage time distributions. By combined study of colloidal dynamics in confinement, transport through a biological pore and the folding kinetics of DNA hairpins, we demonstrate conclusively how a short-time, power-law regime of the first-passage time distribution universally reflects the number of intermediate states associated with each process, irrespective of the lengthscales, timescales or interactions in the system. We thereby establish a powerful method for investigating the underlying mechanisms of complex molecular processes.

---

The concept of an energy landscape is a powerful tool in providing a description of complex natural phenomena. In chemical kinetics, reaction profiles have long been used to qualitatively rationalise the outcomes of chemical reactions, casting light upon preferred mechanisms and the effects of catalysis [1, 2]. In biology, energy landscapes are central to understanding the microscopic origins of processes, including protein folding [3, 4, 5, 6, 7, 8] and selective transport through membrane channels [9, 10, 11, 12, 13, 14]. Elucidating the factors governing the dynamics of stochastic processes may be central to even more diverse problems, such as understanding electron transport [15] or the changing stock prices in financial markets [16, 17, 18]. In spite of this, *quantitatively* resolving the energy landscape that governs an arbitrary process is, in general, very difficult, and requires measurements of transition path times [4]. As such, uncovering energy landscapes represents a fundamental problem in fully understanding complex systems.

The question of how a system explores a known energy landscape has been extensively debated, and multiple computational methods to evaluate the resulting dynamic properties have been proposed [19, 20]. By contrast, the inverse problem – that of determining features of an unknown energy landscape, such as the depth of potential minima or number of intermediate states, from knowledge of the dynamic features of the system – is much more challenging. Recently, however, quantitative links between dynamics and the energy landscape have been proposed from theoretical analysis of complex networks of states [21] where the dynamics were quantified by the first-passage time distribution. Yet to analyse systems with unknown energy landscapes the applicability of such theoretical relationships must be tested experimentally. This can only be achieved with detailed knowledge of both the potential energy landscape and dynamics of the process to allow for the quantitative mapping between these two parame-ters to be probed. Such detailed information is often not available for molecular or nanoscale systems where only certain aspects of the dynamics can be obtained or where the energy landscape is unknown or can be assessed only indirectly. In contrast, mesoscale colloidal model systems represent an ideal test bed for understanding and developing these proposed fundamental connections. Here it is possible to manipulate and control the free energy landscape [22, 23, 24, 25, 26, 27] whilst fully resolving the dynamics with no unknown or hidden degrees of freedom.

In this work, we establish a powerful general method to reveal key details of energy landscapes in experimental systems ranging from the mesoscale to the microscale, through quantitative analysis of first-passage time distributions. We first develop our method by studying the diffusion of colloidal particles in microfluidic channels with controlled potential energy landscapes. Here, we observe characteristic behaviour in the short-time regime of the first-passage time distributions which sensitively reflects the number and depth of potential minima crossed by a particle as it escapes the channel (Fig.1a), consistent with theory [21]. We then demonstrate the wider relevance of our method by analysing the dynamics of, firstly, the chemical ratcheting of a DNA oligonucleotide through a nanoscale pore [28] and, secondly, the folding and unfolding of DNA hairpins [29]. Just as with our colloidal system, we find that in both cases the first-passage time distributions clearly show a power-law regime at short times, from which it is possible to infer the number of intermediates associated with the process. As such, we demonstrate that a purely dynamic measurement of the full first-passage time distribution can be used to uncover quantitative features of an underlying potential energy landscape.

**Figure 1:**
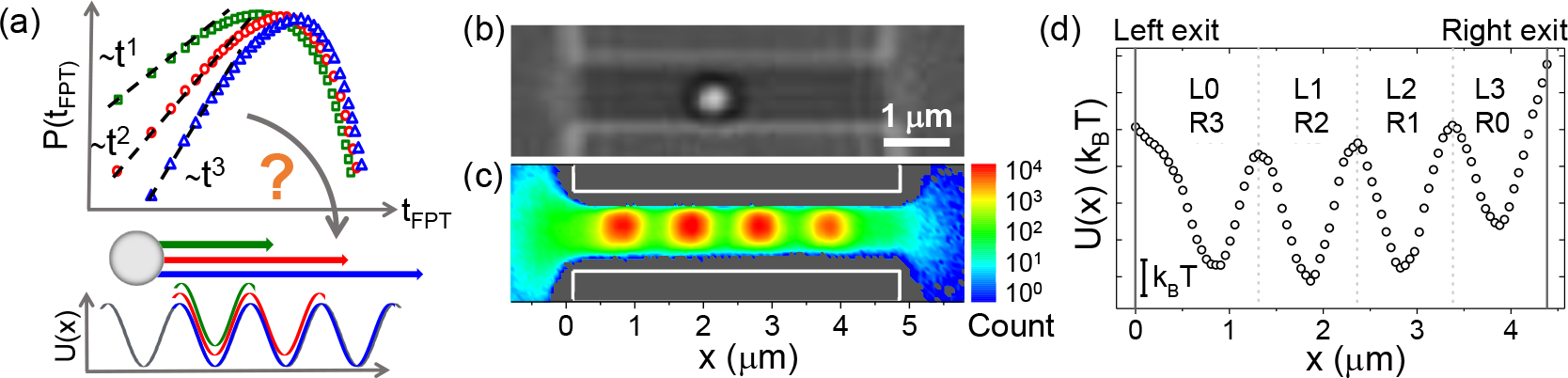
Transport in controlled potential landscapes. (a) An illustration of the proposed link between first-passage time distributions and a particle moving across a potential landscape. A typical experimental image. (c) The experimental 2D probability distribution of particle positions for a channel with four optical traps. The colourbar indicates the total number of times a particle is observed at a position within the channel. (d) The one-dimensional potential landscape, *U*(*x*), calculated from the probability distribution in (c) with potential minima depth Δ*U* ~ 3 *k*_*B*_*T*. Channel exits are indicated by solid lines and the boundaries between minima as dashed lines. Minima labels L(R) 0-3 indicate the lowest number of minima that must be crossed to exit to the left (right) reservoir, for a particle starting in this minimum.

## Establishing first-passage time behaviour in a colloidal system

Colloidal particles are loaded into microfluidic chips designed to include an array of channels linking two three-dimensional reservoirs. Channel dimensions are chosen to confine the colloidal particle to display quasi-one-dimensional diffusion and the sample is imaged using a custom-built, inverted optical microscope (see Fig. 1b). Holographic optical tweezers are used to modulate the potential landscape experienced by the particles by introducing into the channel multiple optical traps. These are designed to be sufficiently weak (depths of 2 to 5 *k*_B_*T*) to allow for escape of the particle from the associated potential minima in observable periods of time. Optical tweezers are also used to automate the data acquisition process [25]. For each first-passage time measurement, a colloidal particle is trapped within the bulk, moved to the centre of the channel, i.e. the boundary between L1 and L2 in Fig. 1(d), released, and allowed to diffuse over the applied potential landscape until it escapes the channel to either the left or right reservoir. Importantly, this automation allows for the acquisition of large datasets (500-4000 trajectories corresponding to more than 10^5^ particle positions). Videos are recorded at 60 Hz, with particle trajectories extracted using standard image analysis techniques.

To quantify the imposed potential landscape we first obtain the joint probability distribution of particle positions, *P*(*x, y*). In Fig. 1(c), *P*(*x, y*) is plotted in 2D for a system with four potential minima of depth Δ*U* ~ 3 *k*_B_*T*. Here, the data clearly show the enhanced probability of a particle residing in the four optical traps. The distribution of particle positions is directly related to the potential of mean force along the *x* axis, *U*(*x*), as *U*(*x*) ~ −*k*_B_*T* ln *P*(*x*), with *P*(*x*) the average over *y* of *P*(*x, y*). This resulting potential landscape for the same four trap system is shown in Fig. 1(d).

For every position in a trajectory, *x*_*i*_(*t*_*i*_), the first-passage time is calculated as *t*_FPT_ = *t*_exit_ − *t*_*i*_, with *t*_exit_ the first time a particle attains a position outside of the channel. As such, while at the start of every measurement the particle is positioned in the center of the channel, the distribution of times to exit from any position within the channel can be measured by assuming the motion of the particle to be Markovian and taking every subsequent position in the trajectory as a new starting point in the analysis. To probe the effect of exit from the channel by crossing an increasing number of intermediate minima, first-passage time distributions are determined for data split into subsets with starting positions, *x*_*i*_, in different regions of the potential landscape (see Fig. 1(d)). Initially separate distributions are obtained for particles exiting to the left (L) or right (R) reservoir before being combined into a single set of distribution functions. For example, in Fig. 1(d), combining data from L1 and R1 would form the full *m* = 1 distribution. The value of *m* indicates the lowest number of minima that must be crossed by the particle, from its initial position (a minima in the channel) to its final position (a reservoir) i.e. *m* = 0 corresponds to crossing no intermediate minima or one boundary in Fig. 1(d), *m* = 1 to crossing one intermediate minimum or two boundaries etc. Typical trajectories for *m* = 0, 1, 2 and 3 are shown in Fig. S1. For all distributions error bars are obtained via the bootstrap method.

Fig. 2(a) and (b) show the first-passage time distributions, *P*(*t*_FPT_), on a linear and log-log scale for colloidal particles diffusing across the potential landscape shown in Fig. 1(d). Distributions for *m* = 1, 2 and 3 are shown, corresponding to the particle starting from a minimum for which exit to a reservoir involves crossing at least 1, 2 or 3 intermediate minima of depth ~ 3 *k*_B_*T* respectively. For clarity, we do not plot *P*(*t*_FPT_) for *m* = 0, which show only the expected exponential decay in all cases. On a linear scale, as *m* increases the distributions exhibit the expected qualitative behavior for diffusion over increasingly large distances, namely a broadening and a shift in the peak to larger times. When plotted on a log-log scale in (b), however, the distributions display a distinct linear behaviour at short-times, with a slope that increases with increasing number of minima that must be crossed. The distributions in Fig. 2(b) are qualitatively different to those of a system with no imposed potential minima (free diffusion), which lack this linear short-time regime (see Fig. S2).

**Figure 2:**
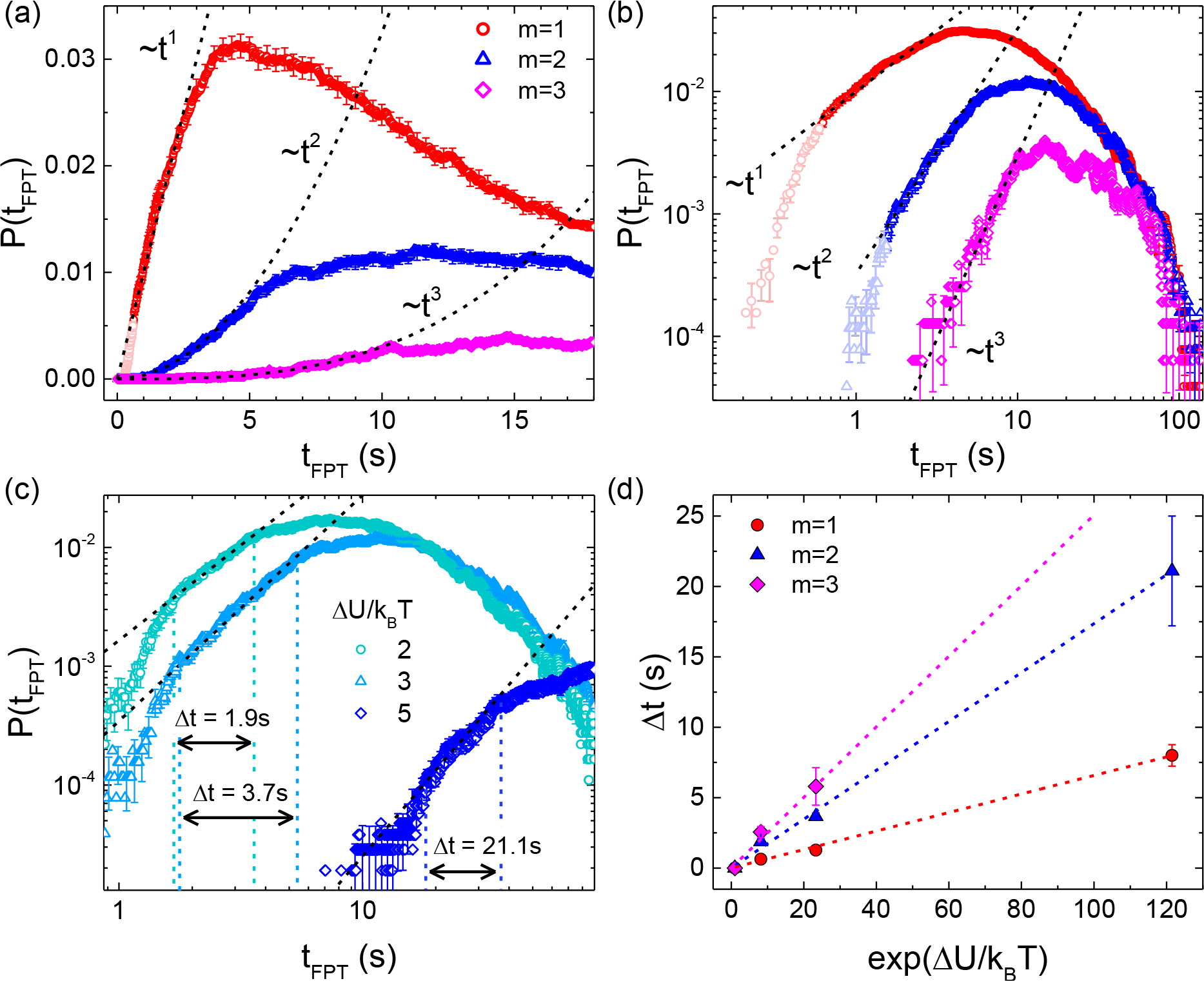
First-passage time distributions of colloidal particles in imposed potential landscapes. Distributions in (a) and (b) show *P*(*t*_FPT_) on a linear and log-log scale for particles diffusing over a potential landscape with minima depth Δ*U* ~ 3 *k*_*B*_*T*. Dashed lines indicate the predicted power-law scaling according to Eq. 1 and the value of *m* for each distribution corresponds to the number of minima crossed by the particle. Panel (c) shows *P*(*t*_*FPT*_) for particles crossing two potential minima (*m* = 2) of different depths, Δ*U*, as indicated. Vertical dashed lines indicate the timescale over which the FPT distributions exhibit a linear regime on a log-log scale, as defined by the black dashed lines. (d) The length of the power-law (linear) regime as a function of potential minima depth for all colloidal systems.

Theoretical results have suggested that the structure of a one-dimensional network of discrete states can be coupled to a short-time scaling of the first-passage time distribution of events starting in a state *A* and finishing in a state *B* (*B* > *A*) via,

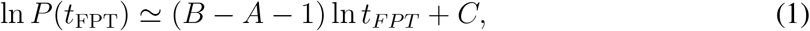

where *C* is a system-dependent constant [21]. This simple quantitative relation is valid because the short-time regime of the distribution for events starting at state *A* and finishing at state *B* is dominated by the shortest trajectories, i.e. those that move directly from *B* to *A* and that do not dwell in any one state for a prolonged time [21]. Remarkably, if a minimum in our continuous landscape is mapped to a ‘state’ in the network description, the linear distortion in the shape of the experimental distributions is exactly consistent with this prediction of a power-law scaling with increasing exponent – on a log scale a linear regime with increasing integer slopes – at short times. To highlight this, dashed lines in Fig. 2(c) and (d) indicate the short-time *t*^*m*^ behavior with, on a log-log scale, the linear regime exhibiting an integer increase in slope with increasing number of minima crossed, exactly as predicted by Eq. 1. We can therefore infer directly from the short-time regime of the first-passage time distribution the number of intermediate potential minima, or equivalently, the number of states, associated with the shortest pathway of the particle to the exit. Note that all experimental trajectories in a dataset are used to build each distribution; trajectories for which the particle has moved back and forth across the landscape simply fall in the long-time regime of the distribution, and so do not effect the slope of the distribution at short times.

At very short-times, however, clear deviations from the linear scaling are seen in *P*(*t*_FPT_) for *m* = 1, 2 (light-coloured points in Fig. 2(b)). This deviation is likely to arise from the fact that the theory assumes discrete states with large barriers whilst the experiment has a continuous potential landscape with depth of a few *k*_B_*T*. To explore this effect further, Fig. 2(c) shows *P*(*t*_FPT_) for *m* = 2 with varying minima depth, Δ*U*, as indicated. On a log-log scale, a linear fit is applied to the linear regime of each distribution and is shown as a dashed black line. In all cases, the slope is close to 2, consistent with the prediction of Eq. 1 for particles that must cross at least two minima to exit the channel. As the potential minima become deeper, however, the linear power-law regime becomes increasingly pronounced and we observe a reduction in the deviation from the linear regime at very short times. A comparison between the data and the linear fit in Fig. 2(c) allows for determination of the length of the linear power-law regime, Δ*t*, as indicated by the vertical dashed lines. In Fig. 2(d) we plot this length of the power-law regime for *m* = 1, 2 and 3 as a function of Δ*U*. For all data, we observe that Δ*t* scales with exp(Δ*U*/*k*_B_*T*) with a gradient that increases with *m*. Fig. 2(d) thus shows that the short-time regime not only provides information on the number of underlying intermediate potential minima, from the slope of the linear regime, but also the depth of the minima, from Δ*t*. This behaviour can be rationalised by considering the time necessary to move across the potential landscape (see supplementary information).

## Inferring molecular intermediates from first-passage times

Turning to nanoscale and molecular systems, we now explore the general applicability of our findings to the more complex dynamic phenomena found at these lengthscales. In particular, we exploit the same analysis to understand two different dynamic processes: firstly, transport in a nanoscale pore – the ‘molecular hopper’ system – and secondly, the folding and unfolding kinetics of DNA hairpins. Details of the molecular hopper system have been reported previously [28]. Specifically, the small-molecule hopper carries a ssDNA cargo along a multi-cysteine track by consecutive thiol-disulfide interchanges. The track is built on a *β* strand facing the lumen of an *α*-hemolysin nanopore. Under an applied potential, the ionic current passing through the pore changes as the ssDNA is ratcheted from one cysteine foothold to another, from which the position of the hopper can be determined (see Fig. 3b).

**Figure 3:**
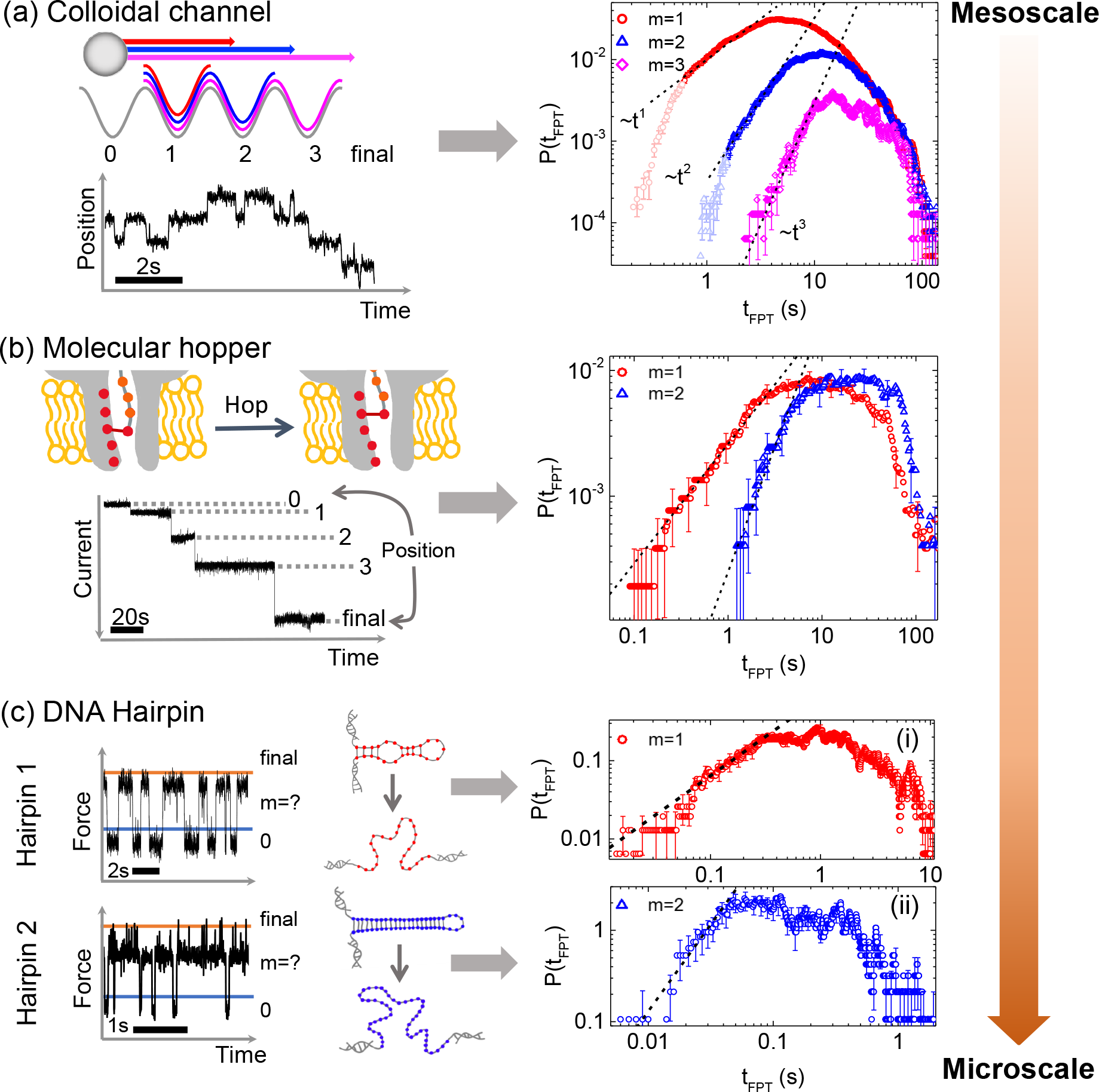
Universal behaviour of the first-passage time distribution from the mesoscale to the microscale. (a) *P*(*t*_FPT_) for the colloidal channel system. (b) *P*(*t*_FPT_) for the nanoscale, ‘molecular hopper’, system comprising a DNA oligonucleotide ratcheted through an *α*-hemolysin pore. *m* = 1 and *m* = 2 distributions are shown for crossing one or two states (making two or three hops, respectively). (c) *P*(*t*_FPT_) for the folding and unfolding dynamics of two DNA hairpin systems with different structures and thus energy landscapes.

To apply Eq. 1 we map each foothold, or possible position of the cargo, to a ‘state’ in the discrete network description. First-passage time distributions are then calculated by identifying regions of the current trace in which the cargo has crossed one (*m* = 1) or two (*m* = 2) intermediate states equivalent to making two or three hops within the pore, respectively. For example, in Fig. 3(b) hopping from position 0 to position 2 involves crossing a single state (position 1) and would thus contribute to the *m* = 1 distribution. All positions within the pore are assumed to be equivalent, such that transitions between any pair of states that are separated by the correct number of intermediate states are combined into a single distribution. From this procedure we obtain 161 realisations of the two-hop *m* = 1 process and 87 realisations of the the three-hop *m* = 2 process. The resulting first-passage time distributions are shown in Fig. 3(b). These distributions clearly exhibit a short-time linear regime on a log-log scale, with a slope very close to that predicted from Eq. 1 for systems crossing one or two states. In contrast to the colloidal experiment, the hopper generally moves in only one direction, demonstrating the applicability of our approach to a system with more directed motion. As such, we note that for the hopper the long time regime of the distribution is made up of trajectories which have dwelled for longer in a relevant state before moving to the exit, rather than those that have moved back and forth. While it is possible for the cargo to make the four hops required for the *m* = 3 distribution, there were only 35 available realisations of this process and it was not possible to resolve this distribution with sufficient accuracy to unambiguously determine the initial slope. The length of the linear regime for the *m* = 2 state is approximately twice that of the *m* = 1 state, which agrees with our findings from the colloidal system. Furthermore, if the much shorter time and lengthscales inherent to the nanopore system are accounted for, the scaling in Fig. 2(d) can be used to obtain Δ*U* ~ 17*k*_B_*T* for the nanopore system (see supplementary information). This is in good agreement with previous estimates [28]. These results demonstrate that our approach is well suited to characterising details of transport in nanopore systems.

Force spectroscopy measurements of DNA hairpin systems have also been previously described [30, 29]. In these experiments, a molecule of interest is tethered to two beads that can be manoeuvred using optical traps to unfold the DNA duplex into single strands. Here, a ‘hopping’ or passive mode protocol is adopted. Changes in the positions of the beads with respect to fixed optical traps are monitored as a function of time. The relative change in bead positions can be directly related to changes in the force exerted upon the bead by the DNA hairpin, with the force exerted varying according to the precise structural configuration of the molecule. Importantly, each structural configuration of the hairpin corresponds to a different minima in the underlying energy landscape for the molecule. From this, the force serves as a proxy for the different structural configurations, linked to different energy minima, and different force values can thus be associated with the different ‘states’ in Eq. 1. Notably, however, direct analysis of the force vs time trace using step fitting is more challenging due to the substantial fluctuations (see Fig. 3(c)), which make it less straightforward to directly determine the number of states and their associated force ranges directly from the trace. These experimental systems thus test the applicability of our approach to molecular systems with lower signal-to-noise ratios, for example, processes that have a well-defined start and end point but where all intermediate states cannot be resolved.

As we can now only consider transitions between the extremes of high and low force, to probe passage over different numbers of states, we consider two different DNA hairpin structures that can adopt different numbers of intermediate configurations (intermediate force states) between the fully folded and unfolded states. To calculate the first-passage time distributions we define a certain force range as the initial state (below the blue line in Fig. 3(c)) and consider the time necessary to first reach a small percentage (< 0.5%) of points in the trace at the opposite extreme of force (indicated by the orange line). The first-passage time distributions calculated in this way for the folding of the two different DNA hairpin molecules are shown in Fig. 3(c). Again, both distributions show a clear linear regime on a log-log scale at short times with (i) exhibiting a slope close to 1 and (ii) a slope close to 2.

Interpretation of the initial scaling of the distributions requires the structure of the two DNA hairpins to be considered. Data used to build the first-passage time distribution plotted in Fig. 3(c)(i) is taken from a short (20bp) DNA hairpin that can transition from a folded (high force) state to an unfolded (low force) state via one intermediate [29]. In transitioning between the folded and unfolded states the system therefore crosses only one intermediate state, i.e. it represents an *m* = 1 process, and this is consistent with our finding that the observed slope in the distribution is equal to one. Data in (ii) is taken from a longer DNA hairpin with a more complex structure and, thus, free energy landscape designed to adopt four different configurational states: an initial configuration, final configuration and two intermediates. As such, in moving from the initial to final configurational state two intermediate states will be crossed, resulting in an *m* = 2 distribution with initial slope equal to two in Fig. 3(c). Remarkably, this demonstrates that our analysis can distinguish between the different energy landscapes associated with the possible configurations of these two different DNA hairpins and demonstrates the ability of our approach to elucidate details of the energy landscape in a molecular system.

## Discussion

We have established a powerful and general protocol that uncovers details of an underlying potential energy landscape by explicitly analysing system dynamics as quantified by first-passage time distributions. Our approach was first rigorously explored in a mesoscale colloidal model system that allows for the resolution of detailed dynamic information in an experimentally controlled potential energy landscape. Here we observe a characteristic scaling in the short-time regime of the distribution that directly reflects the number and depth of potential minima experienced by the particle. This confirms the applicability of theoretical results for a discrete 1D network of states [21] to systems with continuous potential energy landscapes of finite depth.

Having fully detailed the applicability of this approach to our colloidal model system, we broadly applied our method to dynamic processes occurring in nanoscale and molecular systems. More specifically, we calculate the first-passage time distributions for both transport of a molecule through a nanoscale pore [28] and for the folding and unfolding of DNA hairpins [29]. In both cases, the distributions clearly show a power-law regime at short times from which it is possible to infer the number of intermediate minima in energy landscape associated with the process. Hence, our approach can not only be applied to systems from the mesoscale to the microscale, but also to data acquired by a range of different experimental techniques.

The main conclusion of our work is the experimental confirmation across multiple length-scales of a simple, analytic relationship between the dynamics of a process and key underlying aspects of the free-energy landscape that drives it. Consequently, for phenomena where only dynamic information is accessible, calculation of the first-passage time distribution can be reliably used to reveal details of the underlying energy landscape. While we note that out method does not allow for a full restoration of the underlying landscape, the two features obtained from the distributions, namely depth and number of potential minima, are those most important to understanding the dynamics of a process. It is also interesting to note that although in this work only the dynamics in effectively one-dimensional energy profiles have been studied, theoretical predictions exist for multi-dimensional networks of states [21, 31, 32] and it will now be important to explore processes with more complex free-energy landscapes.

## Acknowledgements

A. L. T. and U. F. K. acknowledge Yizhou Tan for help with preliminary colloidal experiments.

## Author Contributions

A. L. T., A. B. K. and U. F. K. conceived the experiments. A. L. T. collected the colloidal data and performed data analysis on all systems. J. G. built the holographic optical tweezers set-up. F. R. and M. R. performed the DNA hairpin experiments, H. B. and Y. Q. performed the molecular hopper experiments. All authors contributed to writing the manuscript.

